# Beneficial effects of cellular coinfection resolve inefficiency in influenza A virus transcription

**DOI:** 10.1101/2022.05.01.490193

**Authors:** Jessica R. Shartouny, Chung-Young Lee, Anice C. Lowen

**Author notes:** These authors contributed equally (Jessica R. Shartouny, Chung-Young Lee).

## Abstract

For diverse viruses, cellular infection with single vs. multiple virions can yield distinct biological outcomes. We previously found that influenza A/guinea fowl/Hong Kong/WF10/99 (H9N2) virus (GFHK99) displays a particularly high reliance on multiple infection in mammalian cells. Here, we sought to uncover the viral processes underlying this phenotype. We found that the need for multiple infection maps amino acid 26K of the viral PA protein. PA 26K suppresses endonuclease activity and viral transcription, specifically within cells infected at low multiplicity. In the context of the higher functioning PA 26E, inhibition of PA using baloxavir acid augments reliance on multiple infection. Together, these data suggest a model in which sub-optimal activity of the GFHK99 endonuclease results in inefficient priming of viral transcription, an insufficiency which can be overcome with the introduction of additional viral templates to the cell. These findings offer rare mechanistic insight into the benefits of viral collective dispersal.

**Graphical Abstract:** 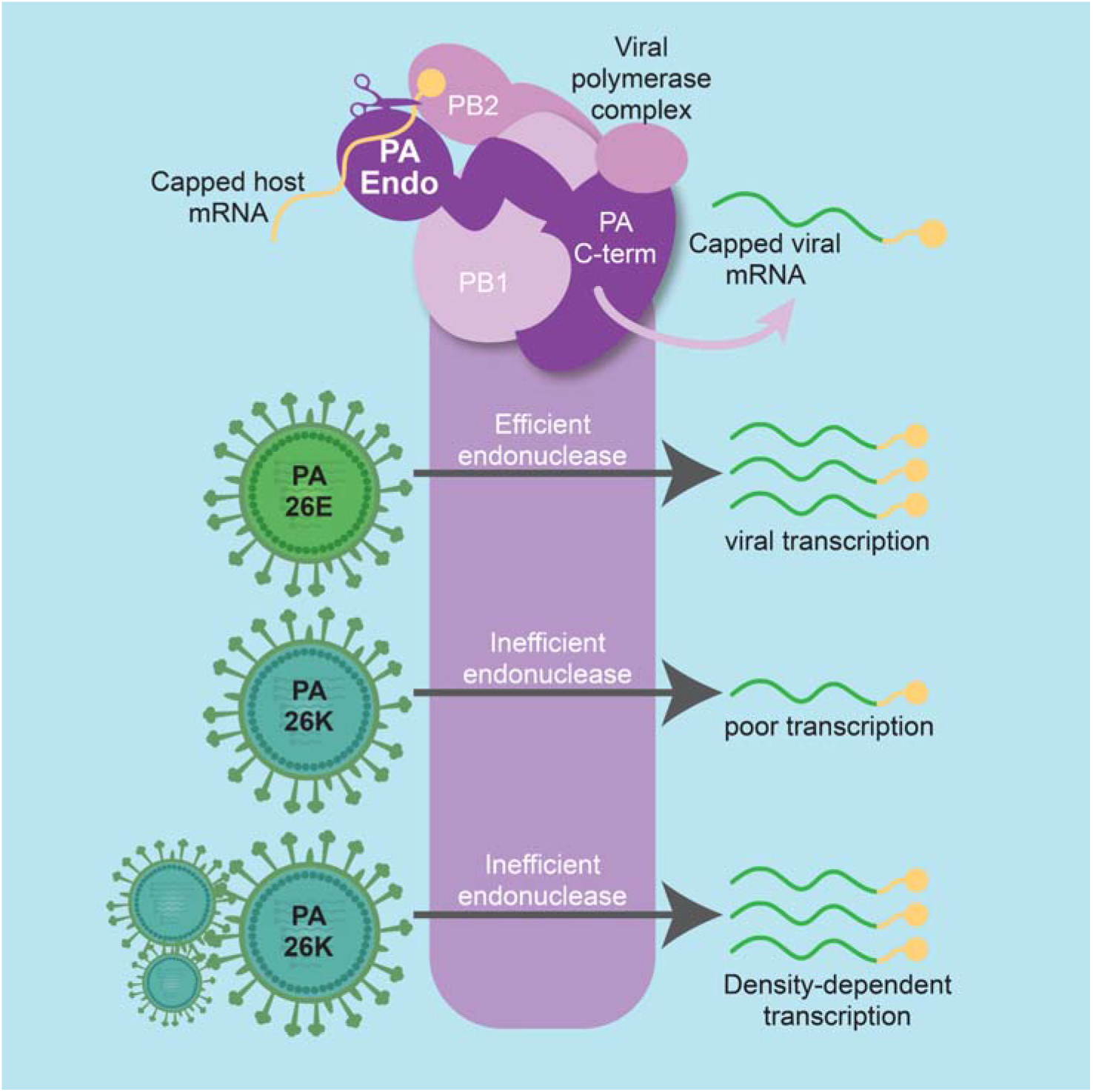

## Introduction

Influenza A viruses (IAVs) impose a substantial burden on public health and agriculture each year. In humans, IAVs circulate seasonally, causing several million illnesses per year globally, and have caused four pandemics since 1918 (Troeger et al., 2019). Fundamental understanding of IAV biology is critical to the design of optimal strategies for prevention and control of influenza.

As has been seen for diverse viral species (Andreu-Moreno and Sanjuan, 2018; Boulle et al., 2016; Chen et al., 2015; Santiana et al., 2018; Zhong et al., 2013) interactions between homologous, coinfecting, IAVs can be highly biologically significant (Brooke, 2017; Jacobs et al., 2019; Phipps et al., 2020). While most single infections are abortive, delivery of multiple viral genomes to a cell strongly increases the likelihood of productive infection and can augment both replication rate and yield (Brooke et al., 2013; Jacobs et al., 2019; Phipps et al., 2020). In other words, the virus-virus interactions that play out during cellular coinfection are typically beneficial and often required for productive infection. Of note, this feature of IAV biology strongly increases the frequency of reassortment, an important source of viral genetic diversity (Steel and Lowen, 2014).

IAV reliance on multiple infection appears to be particularly acute under conditions that are unfavorable for viral replication, such as in a new host species (Phipps et al., 2020). For a G1-lineage strain, influenza A/guinea fowl/Hong Kong/WF10/99 (H9N2) virus (GFHK99, also referred to as WF10), we found that coinfection with multiple homologous viruses was essential for robust replication in mammalian cells, but not avian cells. While every IAV tested thus far has shown some reliance on multiple infection, the strong host-specific reliance of GFHK99 was not apparent for influenza A/mallard/Minnesota/199106/99 (H3N8) virus (MaMN99). Using gene segment reassortments between these two IAVs, we found that the PA segment drives the high reliance of GFHK99 virus on multiple infection. However, the mechanistic basis for this genetic association remained unclear.

The PA gene segment encodes two proteins: PA and PA-X. PA is one of three protein subunits that comprise the viral RNA-dependent RNA polymerase, along with PB2 and PB1. The N-terminus of PA contains an endonuclease that cleaves cellular mRNAs 10-20 bases downstream of the 5’ cap, in a process termed cap-snatching (Dias et al., 2009; Pflug et al., 2017). The resultant capped primer is required for mRNA synthesis by the viral polymerase (Guilligay et al., 2014). PA-X, discovered in the past decade, is produced by ribosomal frame-shifting during translation of the PA mRNA (Jagger et al., 2012). The N-terminal 191 amino acids of PA-X are identical to those of PA, while the C-terminal 41 or 61 amino acids are unique (Gao et al., 2015). PA-X has been shown to contribute to the shutoff of host protein synthesis (Gaucherand et al., 2019).

In this work, we sought to elucidate the mechanistic drivers of the GFHK99 strain’s high reliance on multiple infection, tied to the PA gene segment. Our data demonstrate that the phenotype maps to residue 26K within the PA endonuclease. This amino acid lowers endonuclease activity, leading to inefficient viral transcription in cells infected at low multiplicity of infection (MOI). Conditions conducive to cellular coinfection allow robust transcription by a PA 26K virus in mammalian cells and support enhanced progeny production. Treatment of infected cells with a PA endonuclease inhibitor, baloxovir acid, leads to a similar reliance on multiple infection. These data suggest that cellular coinfection is beneficial because the delivery of multiple copies of the eight viral ribonucleoproteins (vRNPs) to the cell increases the frequency of successful transcription events, allowing infection to be initiated efficiently despite sub-optimal conditions.

## Results

### Reliance on multiple infection of GFHK99 is linked to the endonuclease region of PA

To determine which region of the PA gene segment contributed to the high reliance on multiple infection seen in GFHK99, chimeric PA gene segments were created using GFHK99 PA and MaMN99 PA. The segment was divided into three regions roughly corresponding to the domains of the PA protein (Fig. 1A). Each chimeric segment and the full-length GFHK99 PA were incorporated into the MaMN99 background. To allow virus-virus interactions to be monitored, homologous infection pairs, termed wildtype (WT) and variant (VAR) viruses, were produced for each PA genotype. VAR viruses contained a synonymous mutation in each of the eight gene segments to allow differentiation from WT segments in molecular assays. In addition, WT and VAR virus HA proteins were differentially modified with epitope tags to allow quantification of cellular infection at a protein level (Phipps et al., 2020).

**Figure 1:**
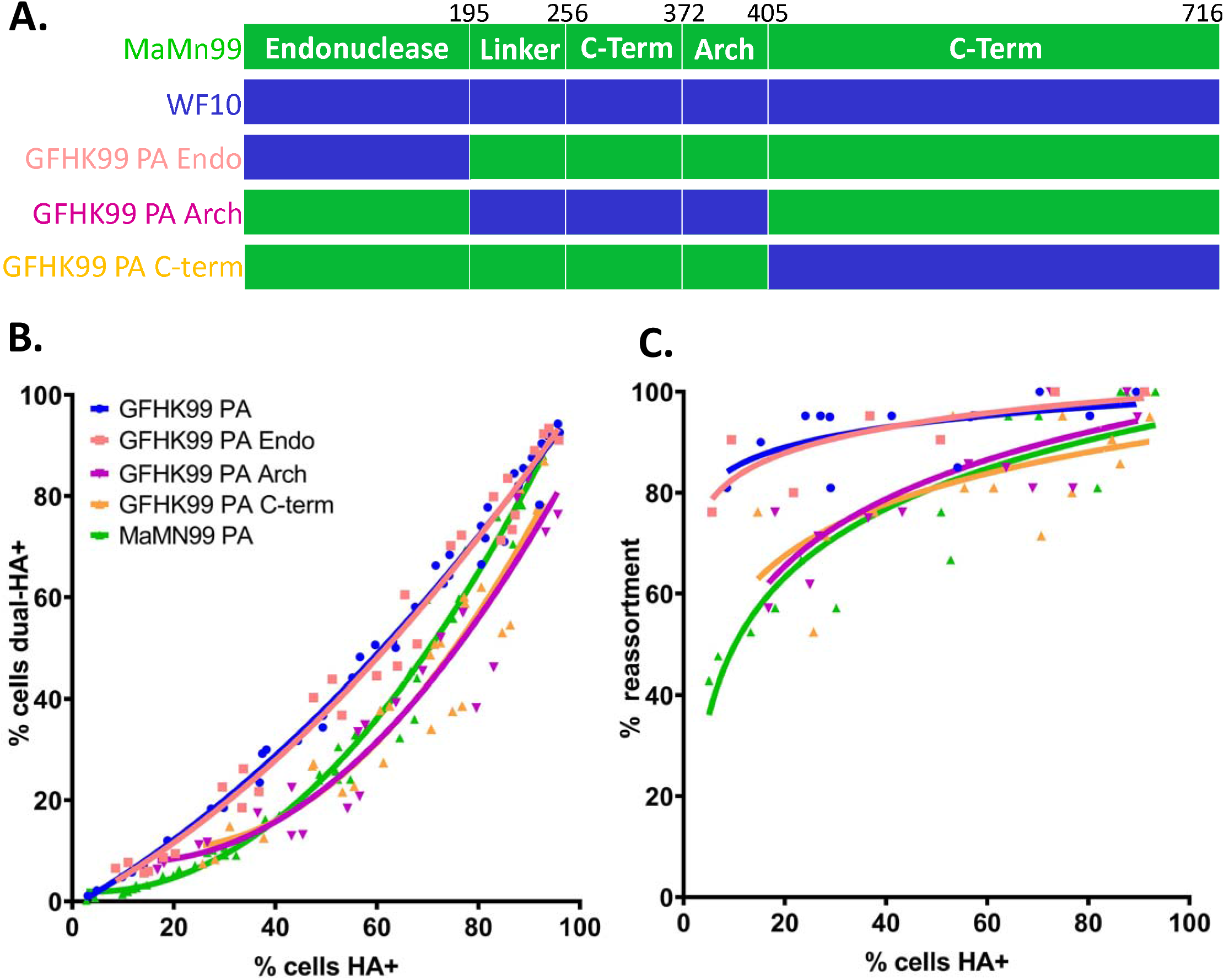
Replacement of the MaMN99 endonuclease region with that of GFHK99 PA increases reliance on multiple infection. **A.** Chimeric PA gene segments that introduce regions of the GFHK99 PA into the MaMN99 PA: the endonuclease (GFHK99 Endo), the middle region (GFHK99 Arch), and the C-terminal domain (GFHK99 C-term). These segments were incorporated into the MaMN99 background. **B.** The relationship between the percentage of the cell population coinfected with WT and VAR (dual HA+) and the percentage that is infected with either or both viruses (HA^+^). Data plotted is from three independent experiments. **C.** The percentage of progeny viruses with any reassortant genotype (% reassortment) is plotted against the percentage of cells expressing hemagglutinin (HA^+^). The PA segment genotype of WT and VAR viruses is indicated in the legend. All viruses carried the remaining seven segments from MaMN99 virus. Data shown for each virus are derived from two independent experiments.

Using flow cytometry, the frequency of cellular coinfection between WT and VAR viruses was evaluated. Since this assay detects viral protein, it measures the extent to which viral gene expression relies on multiple infection. MDCK cells were coinfected with homologous WT and VAR pairs and, to enable quantitative analysis, infections were limited to a single cycle such that progeny viruses cannot be propagated onward. The relationship between the percentage of cells that stained with one or both epitope tags (total HA^+^) and the percentage of cells staining with both tags (dual-HA^+^) was evaluated (Fig 1B). As seen previously, MaMN99 WT and VAR viruses produced distinct populations of singly-infected cells and coinfected cells, with the frequency of coinfection increasing with total infection levels. Conversely, for the GFHK99 PA in a MaMN99 background (GFHK99 PA virus), nearly all infected cells were coinfected with both WT and VAR. Thus, a linear relationship was seen between dual-HA^+^ and total HA^+^, with a slope of 1.02 (95% C.I. 0.979 to 1.07, R^2^= 0.981). Both the GFHK99 Arch PA and GFHK99 C-term PA viruses showed similar infection patterns to that of MaMN99 virus. In contrast, the GFHK99 Endo PA virus displayed a linear relationship between dual-HA^+^ and total HA^+^, like the GFHK99 PA virus. The slope obtained from a linear regression of GFHK99 PA Endo was comparable to that of GFHK99 PA at 1.03 (95% C.I. 0.959 to 1.10, R^2^= 0.971).

The prevalence of reassortant viruses within the progeny virus population was then determined. Since only cells coinfected with WT and VAR viruses can produce reassortants, this assay gives an indication of the relative productivity of singly- and multiply-infected cells. Reassortants were identified by deriving clonal isolates from the progeny population and then genotyping each segment therein. The frequency of reassortants within MaMN99 progeny virus populations was low at lower infection levels and increased as the percentage of HA^+^ cells increased. By comparison, GFHK99 PA virus coinfections resulted in high frequencies of reassortment even at low levels of infection, signifying that most of the progeny viruses were produced in cells infected with both WT and VAR (Fig. 1C). WT-VAR coinfections with GFHK99 Arch and GFHK99 C-term viruses exhibited similar reassortment outcomes as seen with MaMN99 virus. Conversely, GFHK99 Endo PA coinfections displayed high percentages of reassortant progeny, on par with those seen for GFHK99 PA. The high frequencies of reassortant progeny and high levels of dual HA positivity observed in coinfections with GFHK99 Endo PA viruses indicate that the endonuclease region of GFHK99 PA confers a high reliance on multiple infection.

### Disruption of PA-X does not alter reliance on multiple infection

The endonuclease domain is shared by the PA and PA-X proteins. To determine whether PA-X was the driving force behind high reliance on multiple infection, viruses were created in which the PA-X reading frame was disrupted (Gaucherand et al., 2019). Because PA-X is difficult to detect by western blotting, the effectiveness of this disruption was verified at a functional level. Using plasmid transfection, the full length MaMN99 PA gene segment with or without mutations to PA-X was introduced into cells together with a *Renilla* luciferase reporter construct. The wild type PA construct strongly suppressed the reporter signal relative to that seen with the ΔPA-X construct, consistent with the host shut-off activity of PA-X (Gaucherand et al., 2019) and confirming the effectiveness of the mutations introduced (Fig 2A).

**Figure 2:**
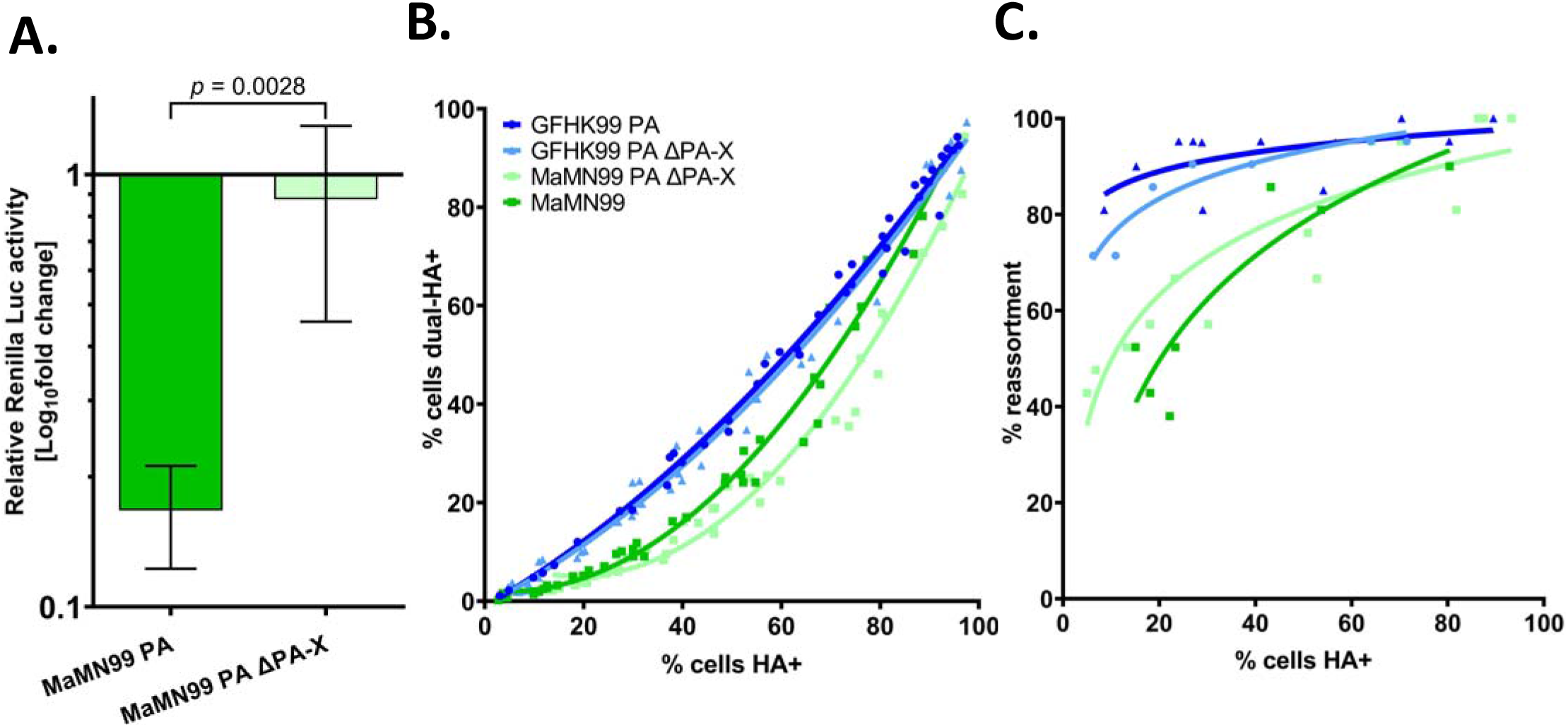
Disrupting PA-X does not alter reliance on multiple infection phenotype of GFHK99 and MaMN99 viruses. **A.** Relative luciferase activity of cells cotransfected with a Renilla luciferase expression plasmid and either MaMN99 PA or MaMN99 PA ΔPA-X expression plasmids. Values represent means ± SDs of three replicates. Statistical significance was analyzed by Unpaired t-test. **B.** The relationship between cells dually-infected with WT and VAR MaMN99 μPA-X or GFHK99 PA μPA-X and cells infected with WT, VAR, or both. Data plotted are from two independent experiments. **C.** The percentage of progeny viruses with any reassortant genotype is plotted against the percentage of cells expressing hemagglutinin (HA). The PA segment genotype of WT and VAR viruses is indicated in the legend. All viruses carried the remaining seven segments from MaMN99 virus. Data shown for each virus are derived from two independent experiments. Results for GFHK99 PA and MaMN99 viruses are reproduced from Figure 1 for comparison.

WT and VAR homologous pairs of these viruses were used to coinfect MDCK cells and the frequencies of coinfected cells and reassortant viruses were examined across a range of MOIs (Fig. 2). By both measures, MaMN99 μPA-X and GFHK99 PA μPA-X viruses displayed similar coinfection reliance phenotypes to their respective parental strain. Since expression of PA-X did not modulate the extent of reliance on multiple infection in either strain, PA-X does not appear to be a major driver of the high reliance phenotype displayed by GFHK99.

### Reliance on multiple infection is dependent on PA 26 in the endonuclease region

An alignment of the PA endonuclease regions of MaMN99 and GFHK99 showed five coding differences at PA amino acids 20, 26, 85, 101, and 118 (Fig. 3A). Owing to the charge difference at PA 26, where MaMN99 has a glutamic acid (E) and GFHK99 has a lysine (K), we focused on this position, which is located proximal to the active site of the endonuclease region (Fig. 3B). Reciprocal mutants were generated in the MaMN99 and GFHK99 PA segments and each was incorporated into the MaMN99 background. WT and VAR homologous pairs of MaMN99 PA E26K and MaMN99: GFHK99 PA K26E (GFHK99 PA K26E) viruses were used in coinfections.

**Figure 3:**
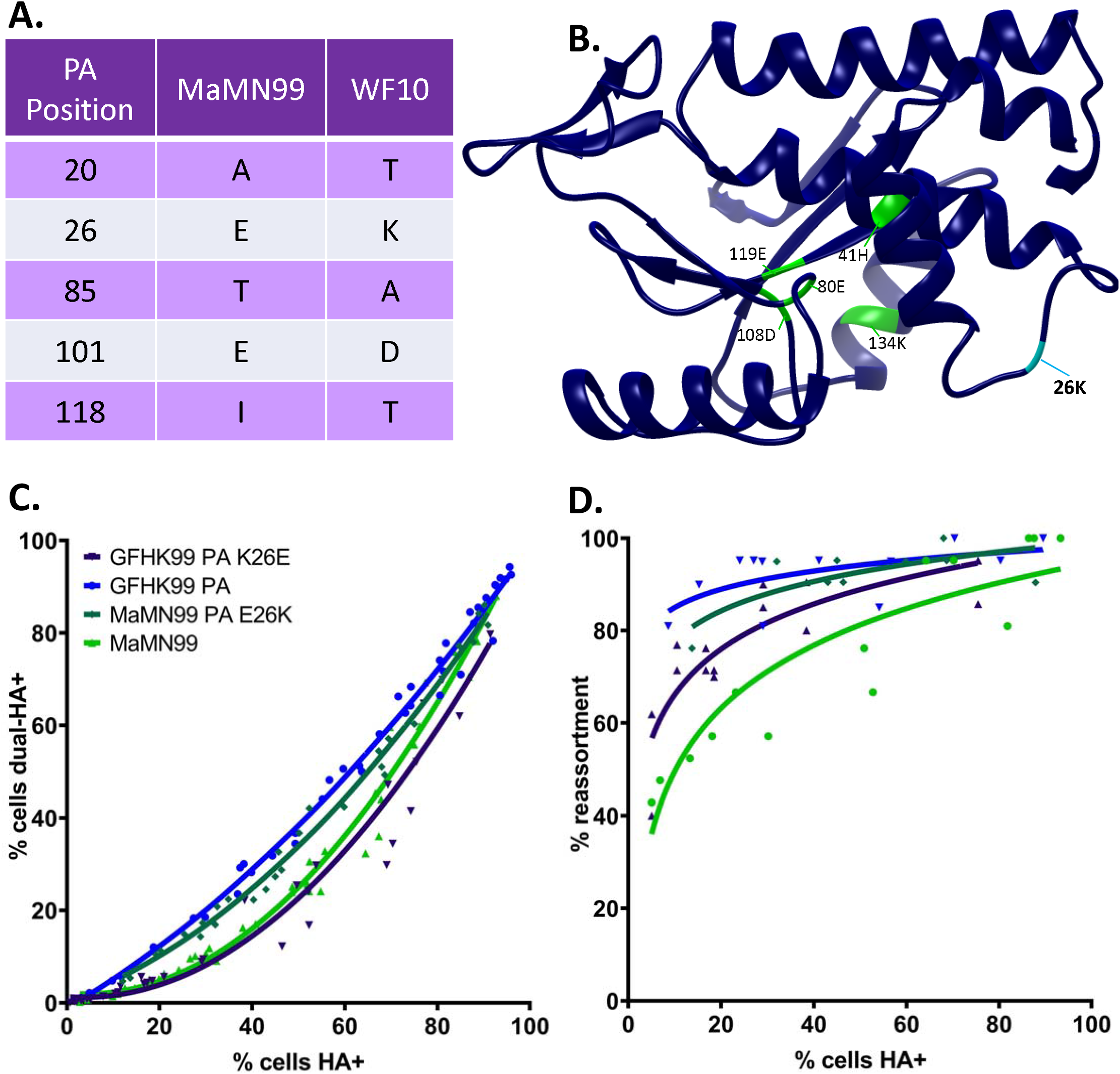
PA 26K is associated with a higher reliance on multiple infection than PA 26E. **A.** An amino acid alignment of MaMN99 and GFHK99 PA endonuclease regions shows five amino acid differences. **B.** The endonuclease domain of GFHK99 PA with the active site residues labeled in green and PA 26K labeled in cyan. **C.** The relationship between the percentage of the cell population infected with both WT and VAR of MaMN99 PA E26K or GFHK99 PA K26E (dual HA+) and the percentage that is infected with either or both viruses (HA+). Data plotted is from three independent experiments. **D.** The percentage of reassortant progeny derived from MaMN99 PA E26K and GFHK99 PA K26E from two independent experiments is plotted against the total percentage of cells expressing HA.

The results showed that swapping the amino acid at PA 26 swapped the phenotypes displayed by MaMN99 and GFHK99 PA. Introduction of K26E within the GFHK99 PA decreased frequencies of dually HA+ cells, while introduction of E26K into the MaMN99 PA had the opposite effect (Fig. 3C). In line with the coinfection results, GFHK99 PA K26E virus yielded fewer reassortants than GFHK99 PA virus, while MaMN99 PA E26K infection progeny were predominantly reassortant (Fig. 3D). Thus, in both PA backgrounds, PA 26K was associated with higher reassortment than PA 26E. Taken together, the data show that viral gene expression and progeny production were focused within coinfected cells to a greater extent for viruses encoding PA 26K compared to those encoding PA 26E.

### Endonuclease activity and transcript production are suppressed by PA 26K

We reasoned that the high reliance on multiple infection resulting from PA 26K might be a result of reduced PA protein levels in infected cells or impeded functionality of the PA protein. To evaluate the first possibility, PA protein levels in cells infected with GFHK99 PA or GFHK99 PA K26E viruses were compared by western blotting (Fig. 4A). GFHK99 PA virus did not display less PA protein than GFHK99 PA K26E virus, indicating that PA 26K did not reduce the accumulation of PA during infection.

**Figure 4:**
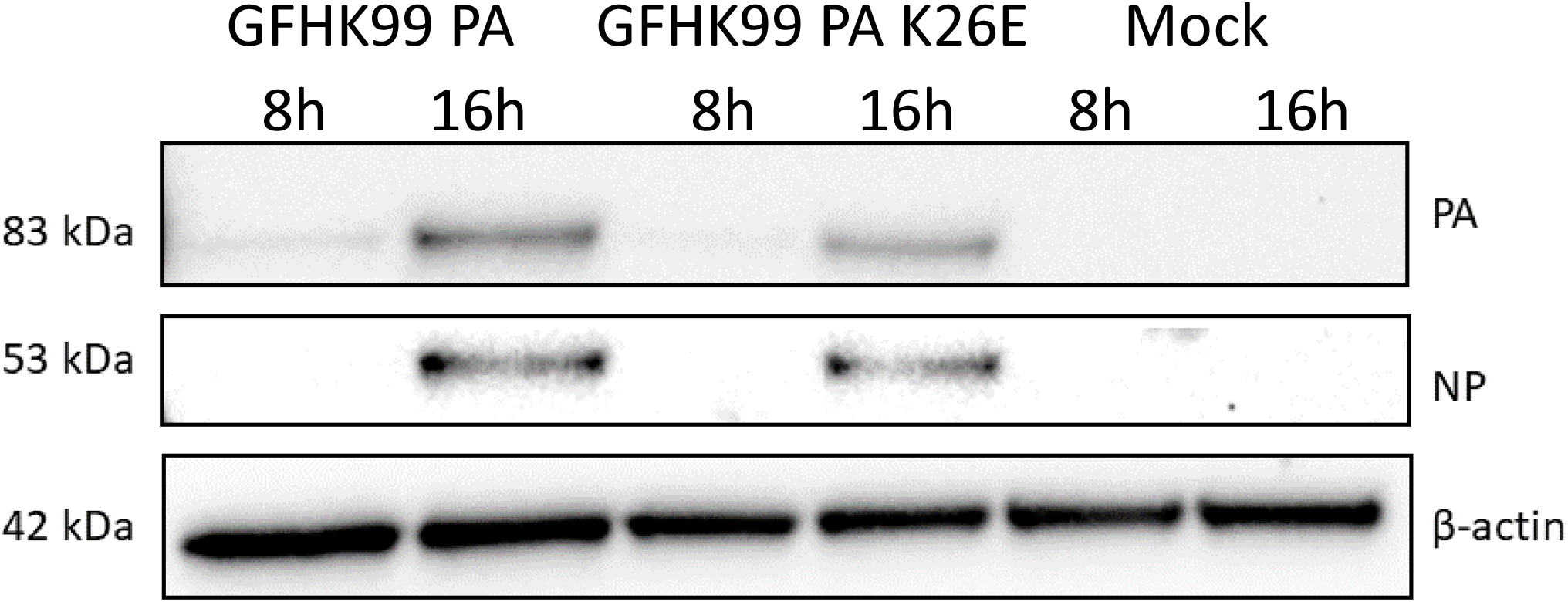
Levels of PA protein are not decreased by PA 26K. Western blot of cell lysates collected at 8 or 16 h post-infection with GFHK99 PA virus, GFHK99 PA K26E virus, or mock infected. Blot was probed for PA, NA, and β-actin.

To evaluate viral endonuclease activity of MaMN99 and GFHK99 PA, we assayed the extent to which the corresponding PA-X proteins disrupted expression of *Renilla* luciferase in transfected cells. This approach was used because PA and PA-X carry the same endonuclease domain, but PA-X activity is more readily monitored in a cell-based assay. As expected based on the mRNA-degrading function of PA-X, the activity of *Renilla* luciferase gradually decreased with an increase in the amount of PA-X plasmid introduced into cells (Fig. 5A). Of note, the effect was markedly weaker with the GFHK99 PA-X than the MaMN99 PA-X, indicating that the GFHK99 PA-X has lower endonuclease activity. Next, E26K or K26E variants of MaMN99 and GFHK99 PA-X, respectively, were tested and PA-X expression of each was verified by Western blot. The E26K mutation in MaMN99 PA-X reduced PA-X activity, showing less reduction of *Renilla* luciferase activity than the wild-type (Fig. 5B, 5D). Conversely, the K26E mutation in GFHK99 PA-X enhanced PA-X activity (Fig. 5C, 5E). Thus, position 26 within the viral endonuclease modulates its enzymatic activity.

**Figure 5:**
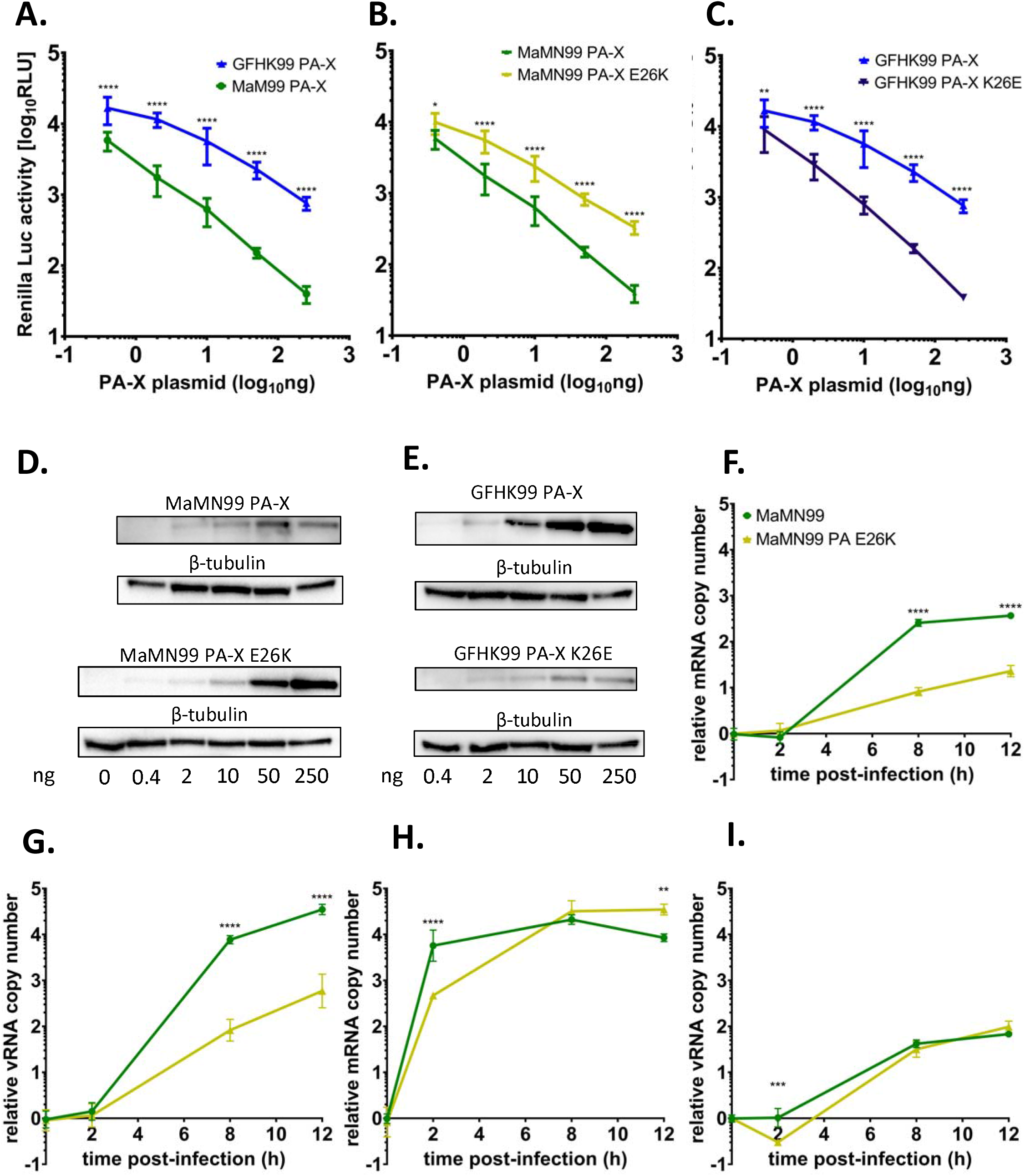
PA 26K confers lower endonuclease activity than PA 26E. A-C. Renilla luciferase expression, plots include three independent experiments, compared via 2-way ANOVA. ****p < 0.0001, *p=0.0201, **p=0.0017. A. MaMN99 PA-X and GFHK99 PA-X. **B.** GFHK99 PA-X and GFHK99 PA-X K26E. C. MaMN99 PA-X and MaMN99 PA-X E26K. **D-E** Western blots show expressed PA-X and β-tubulin at each concentration of plasmid (ng). **F-I.** mRNA (F, H) and vRNA (G, I) were measured at 0, 2, 8, and 12 hpi in a low MOI infection (F,G) and a high MOI infection (H, I). vRNA and mRNA levels were compared via unpaired Student’s t-test.

We further measured the effect of the E26K mutation in MaMN99 PA on viral transcription during infection by measuring the accumulation of viral mRNA under low and high MOI conditions in MDCK cells. At a low MOI, markedly less mRNA was produced in MaMN99-PA-E26K infection compared to MaMN99 infection (Fig. 5F). However, at a high MOI, mRNA accumulated at a similar rate for MaMN99 and MaMN99-PA-E26K viruses (Fig. 5H). These data suggest that low endonuclease activity of PA carrying the 26K polymorphism suppresses viral transcription at a low MOI but is compensated by multiple infection. Measurement of viral genomic RNA (vRNA) in the same cells (Fig. 5G, Fig. 5I) revealed similar patterns, indicating that the effects of PA 26K and multiple infection are also borne out at the level of viral genome replication, most likely as a downstream consequence of the effects on viral transcription.

### Inhibition of endonuclease activity increases reliance on multiple infection

We postulated that inhibiting the PA endonuclease cap-snatching function would enforce an increased reliance on cellular coinfection for productive replication. The drug baloxavir marboxil (Xofluza) targets cap-snatching by chelating the ions in the active site of the PA protein and baloxavir acid (BXA), the active form of the drug, is available for use in cell culture. MaMN99 WT and VAR virus coinfections were treated with an intermediate dose of BXA designed to handicap but not wholly abolish infection (Fig. 6A). The frequency of reassortant progeny resulting from these infections increased across the range of MOIs tested, indicating that nearly all progeny arose from coinfected cells under BXA treatment. This outcome is in stark contrast to that seen in mock-treated MaMN99 infections, where frequencies of reassortment suggest that appreciable levels of virus emanate from singly infected cells.

**Figure 6:**
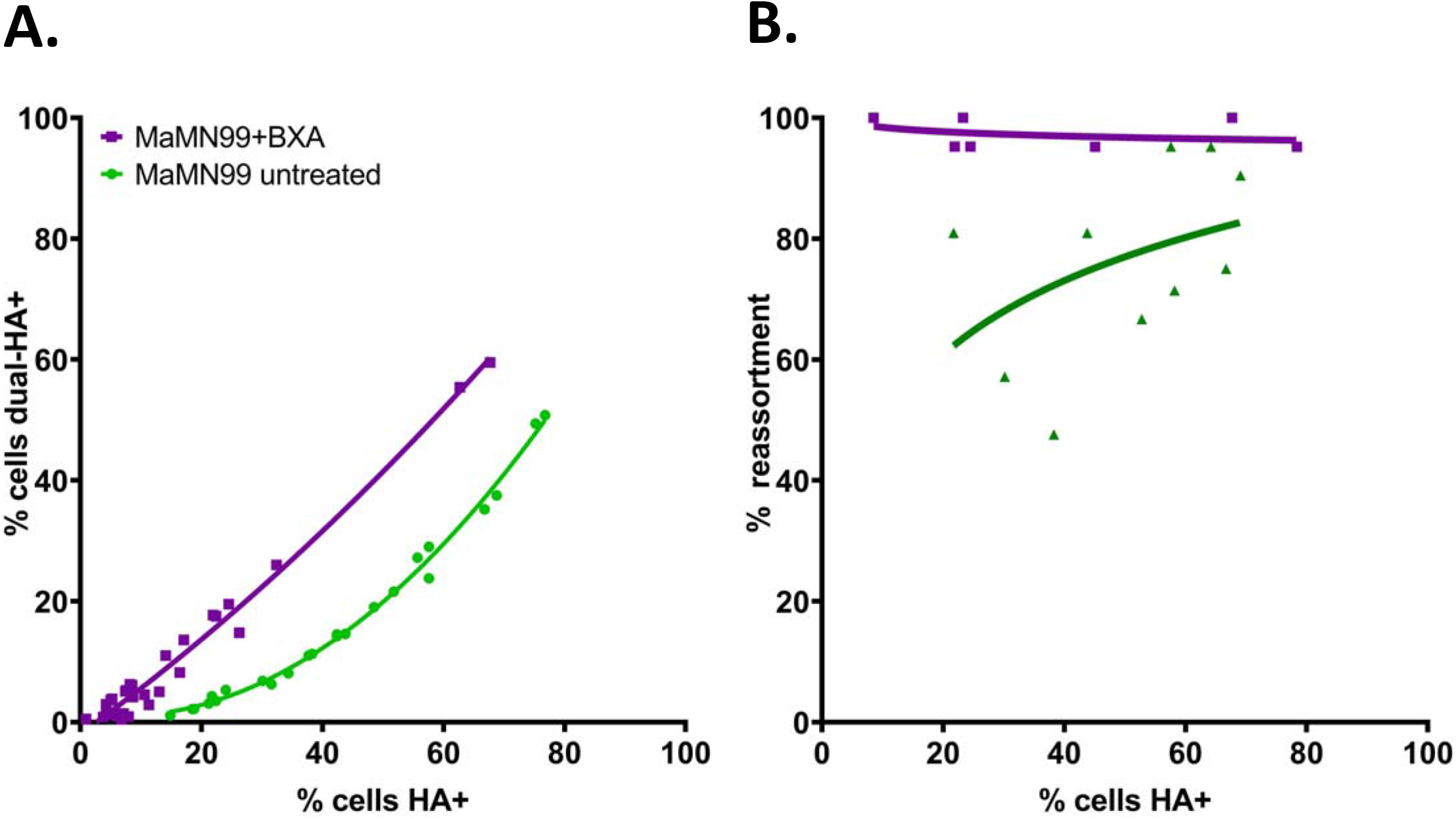
Inhibition of cap-snatching increases reassortment. **A.** The MaMN99 PA endonuclease was inhibited using baloxavir acid (BXA) during a coinfection with homologous WT and VAR viruses and the relationship between percent cells HA+ and cells dual-HA+ was determined for two independent experiments. **B.** The percentage of reassortant progeny resulting from MaMN99 PA +BXA and mock treated was compared to the total percentage of infected cells.

## Discussion

Although cellular coinfection can strongly impact infection outcomes in many virus-host systems, the mechanistic basis for these effects are generally poorly understood. Here we sought to address this deficiency by focusing on GFHK99, an IAV that shows extremely high reliance on cellular coinfection. We identified a polymorphism at position 26 within the PA endonuclease domain as the driver of high reliance on multiple infection. Functionally, PA 26K reduces the activity of the endonuclease enzyme, lowering the efficiency of viral transcription. Importantly, however, under high MOI conditions, the inefficiency of transcription is overcome. Consistent with these results, BXA-imposed inhibition of the PA endonuclease leads to a focusing of viral replication within cells that are multiply-infected. Our data suggest a model in which the delivery of many viral genomes to the cell compensates for inefficient cap-snatching by providing greater opportunity for that process to unfold. The result is a positive density dependence, in which per capita viral reproduction increases as more viruses infect the same cell.

To successfully propagate its genome, an infecting virus must complete a number of discrete steps of the early life cycle, many of which are the targets of cellular defense mechanisms. For IAV, each barrier must be overcome by eight unique vRNP complexes. This process appears to be prone to failure, such that the likelihood of a single IAV establishing an infection is extremely low (Brooke et al., 2013; Heldt et al., 2015). Consistent with low rates of productive infection, IAV infected cells typically support expression from and replication of fewer than eight segments (Brooke et al., 2013; Jacobs et al., 2019). In the case of GFHK99 virus, incomplete viral genomes were detected with a frequency of approximately 90% (Phipps et al., 2020). While high, this frequency is not sufficient to account for the extremely high reliance on multiple infection exhibited by this strain in mammalian systems (Phipps et al., 2020). The data reported here indicate that this additional reliance stems from a deficiency in viral cap-snatching, the process by which the IAV polymerase acquires capped RNA oligomers to prime transcription (Dias et al., 2009). With multiple viral genomes infecting together, the larger number of templates (and associated polymerases) available for primary transcription appears to enable levels of mRNA synthesis sufficient to sustain robust infection.

Our previous work revealed that the extent to which GFHK99 virus relies on multiple infection for productive infection is strongly dependent on host species: in avian systems, the need for multiple infection was greatly diminished (Phipps et al., 2020). Since we focused our studies herein on mammalian systems, it is not clear whether the GFHK99 PA enzyme is more active in avian cells or whether this avian virus is simply less sensitive to the cap-snatching deficiency in a host environment to which it is well adapted. This is a question to pursue in future work.

In using BXA treatment to substantiate the link between deficient cap-snatching and the need for many incoming viral genomes, we show that - within multiply-infected cells - viral replication can proceed under treatment with the active form of an FDA-approved antiviral drug. While potentially consequential for clinical use of baloxavir marboxil and the development of resistance (Hayden et al., 2018; Omoto et al., 2018), it is important to consider that increasing the dose of BXA in our experimental system led to a complete block of infection. In fact, our data more broadly suggest that the targeting of viral cap-snatching and other steps of the viral life cycle that precede genome replication are likely to be efficient therapeutic strategies: this approach is expected to increase the frequency of abortive infections, potentially pushing the rate of productive infections within a host below a threshold needed to sustain viral propagation.

The critical mutation identified, PA 26K, occurs rarely in sequenced influenza strains, including those of the G1 lineage of H9N2 viruses (Clements et al., 2020). Deleterious effects of PA 26K in mammalian and, to a lesser extent, avian systems were identified in previous work using the GFHK99 strain (Clements et al., 2020). PA K26E was also noted as a common variant arising within quail experimentally inoculated with GFHK99 (Obadan et al., 2019). While we see herein that the fitness effects of PA 26K are ameliorated in the context of multiple-infection, these fitness effects are nonetheless likely to explain its low prevalence in nature. We speculate that the occurrence of PA 26K in GFHK99 is the result of genetic drift, which can lead to fixation of deleterious mutations. While PA 26K is not common in circulating IAVs, the phenotype it confers gives an opportunity to identify viral and cellular processes that can underlie beneficial effects of multiple infection.

Cellular coinfection modulates the biology of diverse viruses. In VSV, inducing multiple infections via virion aggregation can accelerate viral production such that it outpaces innate antiviral responses (Andreu-Moreno and Sanjuan, 2018). Similarly, in HIV-1, delivery of multiple viral genomes through cell-to-cell transmission of infection leads to more rapid onset of viral gene expression (Boulle et al., 2016; Zhong et al., 2013). For rotavirus and norovirus, vesicle-bound packets of viral particles have enhanced infectivity relative to free virions and are important vehicles for transmission (Chen et al., 2015; Santiana et al., 2018). Importantly for IAV, beneficial effects of multiple infection can enable viral replication in novel host species (Phipps et al., 2020).

This work reveals one mechanism by which multiple infection can be beneficial for IAV replication; we predict that many distinct mechanisms will lead to a similar effect and that these mechanisms will typically be active at the earliest stages of the viral life cycle, before incoming vRNAs are replicated. Our data underline the potential for multiple infection to enable viral replication under adverse conditions, such as in the presence of deleterious mutation or under antiviral drug treatment. They furthermore reveal mechanistic insight into the high levels of reassortment that are a major feature of IAV dynamics within hosts (Ganti et al., 2021; Tao et al., 2014).

## Author Contributions

Concept and experimental planning was performed by JRS, CYL, and ACL. Data was collected and analyzed by JRS and CYL. Manuscript and figures were written, designed, and edited by JRS, CYL, and ACL. Research funding was acquired by ACL.

## Acknowledgements

We thank Lucas Ferreri for helpful discussion and critical reading of the manuscript. This research project was supported in part by the Emory University School of Medicine Flow Cytometry Core. The work was supported by funding from NIH/NIAID under the Centers of Excellence for Influenza Research and Response contract no. 75N93021C00017 and R01 AI127799.

## STAR Methods

### Lead Contact

Further information and requests for resources and reagents should be directed to and will be fulfilled by the lead contact, Anice Lowen.

### Materials Availability

Reagents are available by request to the lead contact.

#### Data and code availability

- Data generated in this study will be publicly available on GitHub on the date of publication.
- This paper does not report original code.
- Any additional information required to reanalyze the data reported in this paper is available from the lead contact upon request.

#### Method Details

##### Cells and cell culture

Madin-Darby canine kidney (MDCK) cells and 293T cells (ATCC, CRL-3216) were maintained in minimal essential medium supplemented with 10% fetal bovine serum and 100 ug/mL normocin. MDCK cells gifted by Peter Palese, Icahn School of Medicine at Mount Sinai were used for experiments. MDCK cells gifted by Daniel Perez, University of Georgia, were used for plaque assays. All cells were maintained at 37°C and 5% CO_2_ in a humidified incubator and monitored monthly for mycoplasma contamination.

##### Viruses

Influenza A viruses used in these experiments were generated via reverse genetics (Fodor et al., 2000; Neumann et al., 1999). Briefly, 293T cells transfected with ambisense plasmids encoding the eight viral gene segments were injected 16-24 h after transfection into the allantoic cavity of 10-11 day-old embryonated chicken eggs (Hyline International) and incubated for 32-36 h at 37°C. Allantoic fluid was collected and used as the virus stock for experiments. Infectious titers were determined by plaque assay in MDCK cells and by flow cytometry targeting virally encoded epitope tags. Levels of internally deleted defective interfering segments derived from PB2, PB1, PA, and NP segments were confirmed to be minimal for each virus stock, using previously described procedures (Schwartz and Lowen, 2016). All viruses used contained PB2, PB1, HA, NP, NA, M, and NS segments derived from influenza A/mallard/Minnesota/199106/99 (H3N8) virus (MaMN99). MaMN99 PA viruses contained the PA gene segment from MaMN99. MaMN99: GFHK99 PA viruses contained the PA gene segment from influenza A/guinea fowl/Hong Kong/WF10/99 (H9N2) virus. The HA segment of each virus used was engineered to contain either a 6xHIS or an HA epitope tag and GGGS linker following the signal peptide. Homologous WT and VAR viruses were tagged with opposite epitopes to allow differentiation of infected cells via flow cytometry. GFHK99 Endo PA and MaMN99 viruses were tagged WT as HIS-tag and VAR as HA-tag. All other viruses were tagged WT as HA-tag and VAR as HIS-tag. Homologous VAR viruses included one synonymous mutation in each segment relative to the WT strain, as detailed previously (Phipps et al., 2020).

##### Generation of modified PA plasmids

Plasmids encoding chimeric PA gene segments were created using the NEBuilder HiFi DNA Assembly Master Mix (New England Biosciences) according to the manufacturer’s instructions. PCR-amplified fragments encoding the endonuclease region (GFHK99 Endo), linker and arch region (GFHK99 Arch), or C-terminal region (GFHK99 C-term) of GFHK99 PA were combined with a fragment containing the rest of the MaMN99 PA segment in a pDP2002 plasmid, a gift from Daniel Perez (Perez et al., 2017). Mutations to the PA 26 codon were introduced using site-directed mutagenesis (QuikChange, Agilent). Two nucleotide changes were introduced to avoid reversion. MaMN99 ΔPA-X and MaMN99:GFHK99 PA ΔPA-X were designed as described previously (Gaucherand et al., 2019): site-directed mutagenesis of three bases at the X-ORF (t597c, t600c, and t627a) decrease the likelihood of frameshifting and add a TAG stop codon to truncate the C-terminal region of PA-X. Sequences of purified plasmid preparations were verified by Sanger sequencing (Genewiz). WT and VAR pairs of each recombinant virus were generated as described above. ΔPA-X segment loss-of-function was verified via host shutoff capacity: 293T cells were transfected with 40 ng of either pCAGGS-MaMN99-PA or pCAGGS-MaMN99-PA-APA-X plasmids and 50 ng of pRL-TK plasmid (Promega). At 24 h post-transfection, the transfected cells were lysed and 20 μl of lysate was transferred to a 96-well plate. 100 μl of *Renilla* luciferase assay reagent (Promega) was added and then *Renilla* luciferase activity was measured on a Synergy H1 Hybrid Reader (BioTek). *Renilla* luciferase activity was plotted relative to empty vector transfected cells.

##### Infection of cells for quantification of cellular coinfection and viral reassortment

Infections of cultured cells were performed as described previously (Marshall et al., 2013; Phipps et al., 2020). Briefly, homologous WT and VAR viruses were mixed in equivalent amounts based on infectious titers as determined by flow cytometry, diluted serially in 1x PBS, and used to inoculate 80% confluent MDCK cells in 6 well dishes. Synchronized single-cycle infection conditions were used: to synchronize viral entry, virus was allowed to attach during a 45 min incubation at 4°C before addition of warm virus medium (1xMEM, 4.3% BSA, 100 IU penicillin/streptomycin) and incubation for 2 h at 37°C. At the end of this 2 h incubation, residual inoculum was inactivated using a 5 min acid wash in PBS-HCl (pH=3). Cells were then placed in virus medium supplemented with 20 mM NH_4_Cl and 50 mM HEPES and incubated at 37°C. This high pH buffer prevents endosomal acidification and therefore blocks any further viral entry, imposing single cycle conditions. Released virus and cells were collected at 16 hpi (with 0 hpi defined as the time of warming). Supernatant was stored at −80°C until use in plaque assays.

Coinfections with baloxavir acid (MedChemExpress, CAS No. 1985605-59-1) were completed in the same manner as above, with 5nM BXA added to the virus medium both during the 2 h viral entry period and the 14 h viral replication period.

##### Quantification of infection and coinfection

Frequencies of infection and coinfection in cell monolayers co-inoculated with WT and VAR viruses were evaluated based on surface expression of HA- and HIS-tags. Samples were stained for 45 min on ice with Penta HIS Alexa Fluor 647 conjugated antibody (5 ug/ml; Qiagen) and Anti-HA-FITC Clone HA-7 (7 ug/ml; Sigma Aldrich). Cells were then washed and resuspended in PBS-2% FBS in cluster tubes for flow cytometry analysis on a BD-FACSymphony A3 cytometer in the Emory University Flow Cytometry Core. Analysis was performed using FlowJo software. Non-linear regressions (least squares) and linear regressions of the data were performed in Prism Graphpad.

##### Quantification of reassortment

The frequency of reassortant viruses was determined as described previously (Ganti et al., 2021). Plaque assays were performed in 10 cm-diameter dishes to isolate viral clones and agar plugs were collected with 1 mL serological pipettes into 160 μl PBS. vRNA was extracted using the Zymo 96 viral RNA extraction kit then reverse transcribed using Maxima reverse transcriptase (Thermofisher) per the manufacturer’s instructions. cDNA was diluted 1:4 in nuclease-free water and combined with segment-specific primers (Phipps et al., 2020) to differentiate WT and VAR segments by high-resolution melt analysis with Precision Melt Supermix (Bio-Rad) using a CFX384 Touch Real-time PCR detection system (Bio-Rad). Data were analyzed using Precision Melt Analysis software (Bio-Rad) to assign a genotype based on the combination of WT and VAR segments in each isolate. Percent reassortment was calculated as the number of viral isolates with any reassortant genotype divided by the number of isolates screened, multiplied by 100. Results were plotted as a function of percent HA+ cells as determined by flow cytometry. Semi-log curves were fitted to the data in Graphpad.

##### Analysis of the shutoff of cellular gene expression

250, 50, 10, 2, or 0.4 ng of MaMN99 or MaMN99 PA E26K plasmids were ectopically transfected to 293T cells with 50 ng of pRL-TK plasmid using X-tremeGENE 9 (Roche). At 24 h, the transfected cells were lysed and 20 μl of lysate was transferred to a 96-well plate. 100 μl of *Renilla* luciferase assay reagent (Promega) was added and then *Renilla* luciferase activity was measured on a Synergy H1 Hybrid Reader (BioTek).

##### Strand-specific quantification of vRNA and mRNA over time

MDCK cells (1 x 10^5^ cells per well) were seeded onto 24-well plate and incubated at 37°C for 24 h. The cells were washed three times with PBS, then chilled viruses were inoculated under single cycle infection conditions, low MOI of 0.5 RNA copies/cell and high MOI of 1000 RNA copies/cell. After 1h of absorption, the cells were washed three times with PBS. The cells were collected at 0, 2, 8, and 12 h post-infection, and the RNA was extracted using RNeasy Mini kit (Qiagen). To reverse-transcribe specific RNA species, two different primers targeting vRNA or mRNA of NA segment were used (Phipps et al., 2020). The quantitative PCR was conducted with Ssofast EvaGreen Supermix (Bio-rad) and specific primer set targeting vRNA or mRNA of NS using CFX384 Touch Real-time PCR (Bio-rad)(Kawakami et al., 2011).

##### Western blotting

Western blotting was performed using the Thermo scientific miniblot module. SDS-PAGE of reduced samples was run on Bolt Bis-Tris 4-12% premade gels in MOPS. Blotting was performed onto 0.2nm nitrocellulose membranes at 15V for 30 minutes. Blots were blocked with 5% milk in PBS-0.05% Tween 20 for 1h at room temperature, then incubated overnight at 4°C with primary antibodies at 1:2,000: rabbit anti-PA (Genetex), mouse anti-β-actin (Sigma, clone AC-74), and mouse anti-influenza NP (Kerafast, clone HT103). Secondary antibodies (1:3,000) anti-Rabbit IgG-HRP and anti-mouse IgG HRP were incubated with the blots for 1h at room temperature and the blots were developed using Bio-Rad Clarity Western ECL Substrate. Images were taken on the Bio-Rad ChemiDoc MP imaging system.

##### Protein modeling

Models of PA were created using the Phyre2 protein fold recognition server to predict structures based on the published sequence of GFHK99 PA (Genbank accession # MN267497.1) (Kelley et al., 2015). Visualization was performed with UCSF Chimera, developed by the Resource for Biocomputing, Visualization, and Informatics at the University of California, San Francisco, with support from NIH P41-GM103311 (Pettersen et al., 2004).

##### Quantification and statistical analysis

Analysis of these data was performed using the GraphPad Prism statistical software. Replicate sizes are indicated on figure legends where applicable. Linear and non-linear (least-fit) regressions were performed on data collected via flow cytometry. Semi-log lines were fitted to reassortment data. PA-X host shutoff capacity was analyzed using unpaired Student’s t-tests and reported +/− SD for Fig. 2A. Data in Fig. 5 were analyzed via 2-way ANOVA. Alpha = 0.05.

## KEY RESOURCES TABLE

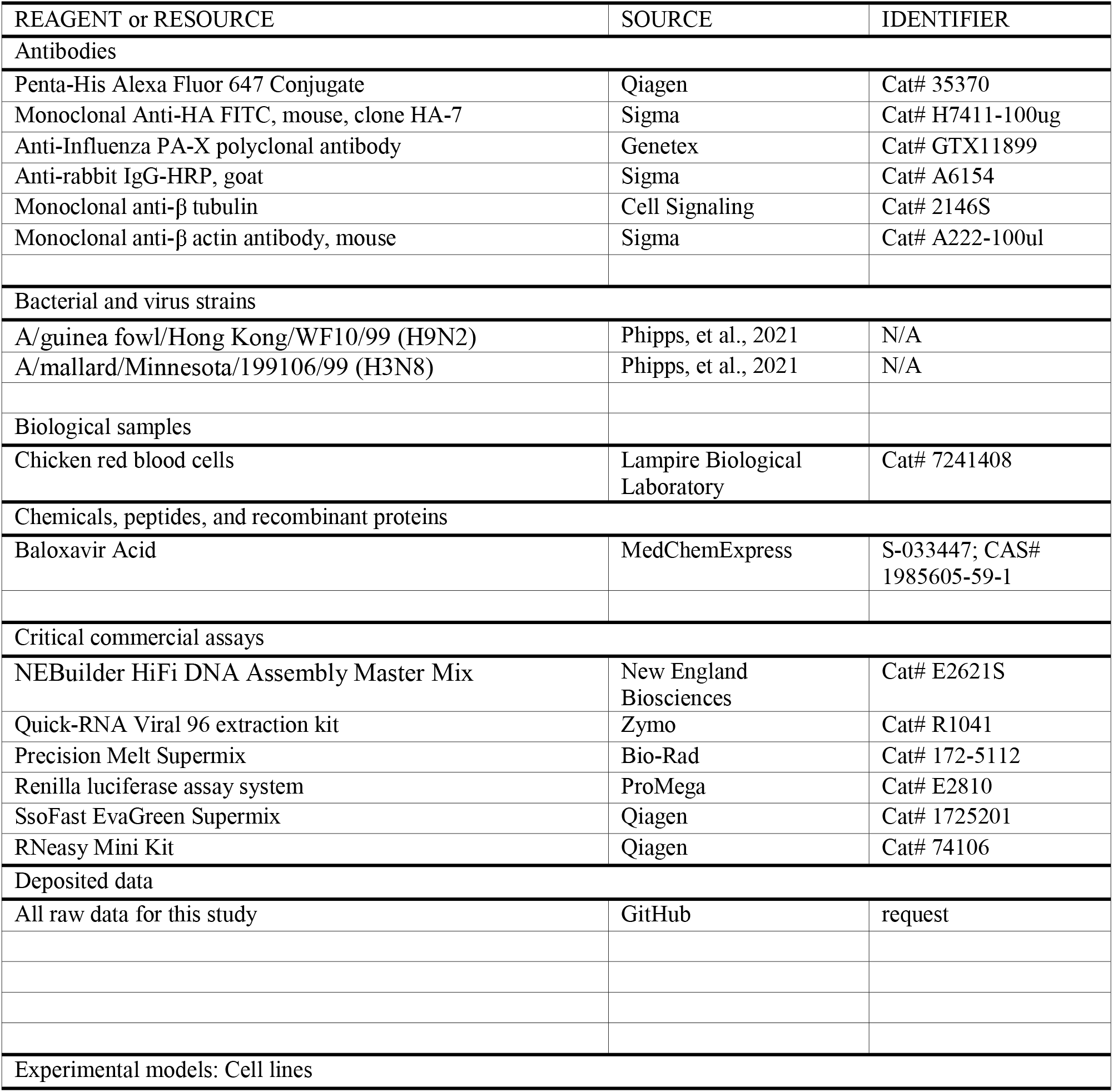
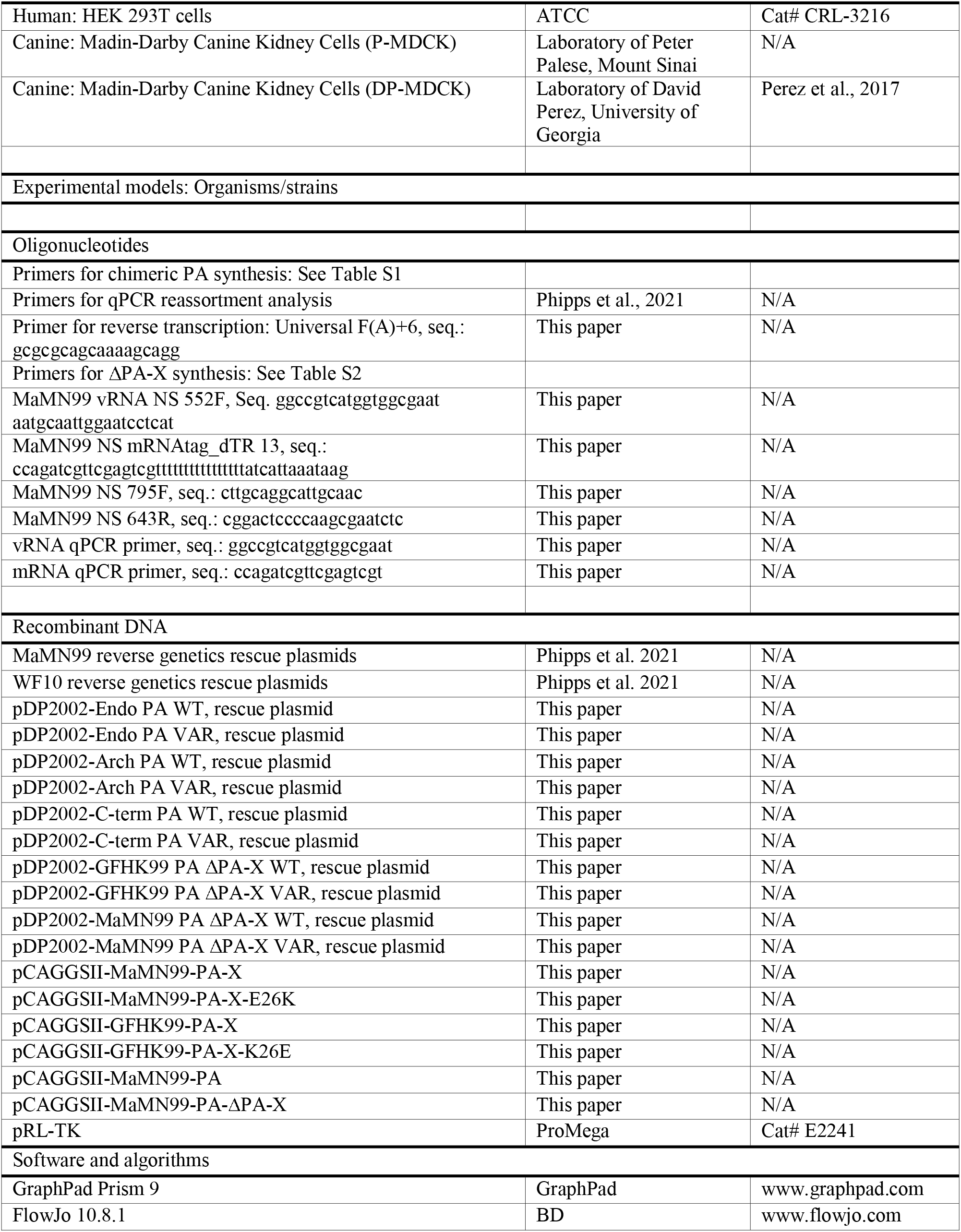
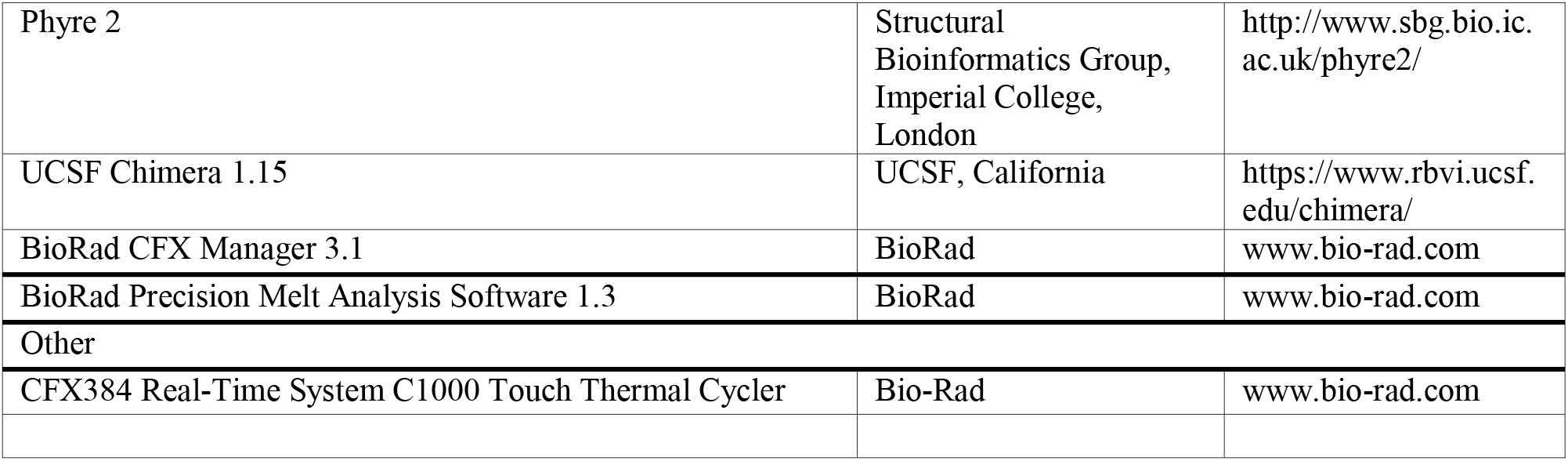

## Supplemental tables

**Table S1:**
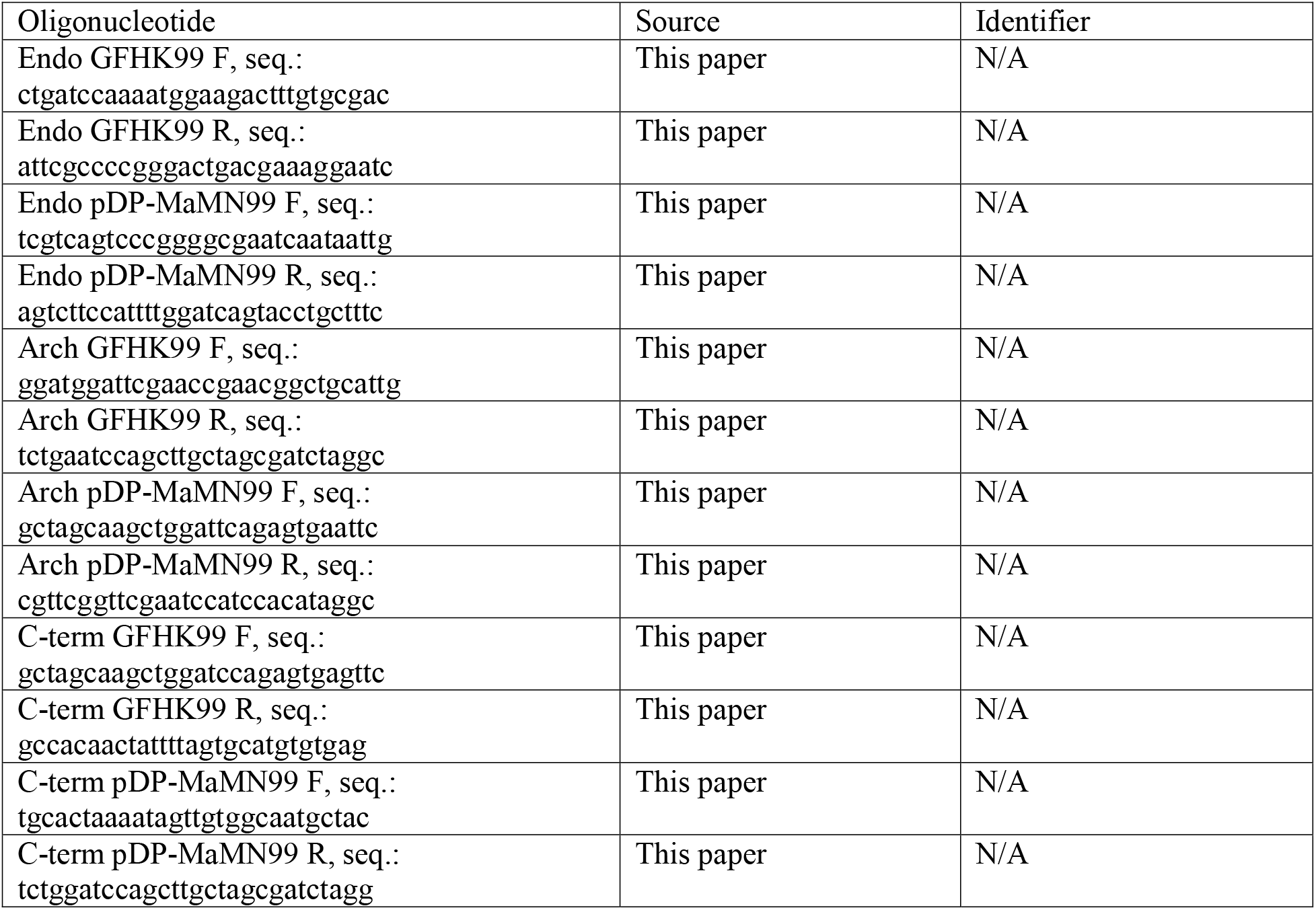
STAR Methods oligonucleotide list, primers used for chimeric PA segment Gibson Assembly

**Table S2:**
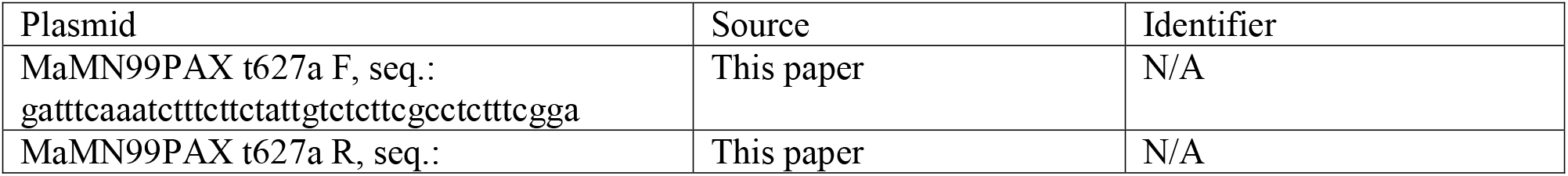
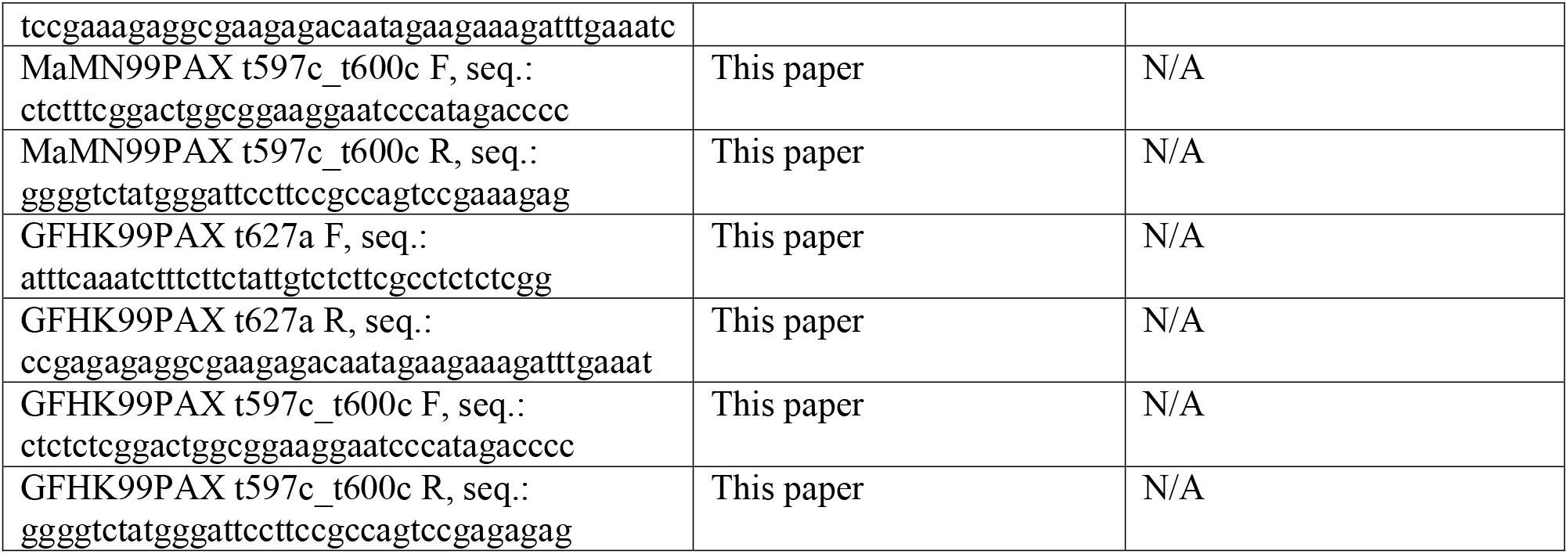
STAR Methods primers for ΔPA-X site-directed mutagenesis

